# Effective target genes for RNA interference-based management of the cabbage stem flea beetle

**DOI:** 10.1101/2024.04.30.591975

**Authors:** Doga Cedden, Gözde Güney, Xavier Debaisieux, Stefan Scholten, Michael Rostás, Gregor Bucher

## Abstract

The cabbage stem flea beetle (CSFB, *Psylliodes chrysocephala*) is a key pest of oilseed rape. The ban on neonicotinoids in the European Union due to environmental concerns and the emergence of pyrethroid-resistant populations have made the control of CSFB extremely challenging. In search of a solution, we have recently shown that RNA interference (RNAi) has potential in the management of CSFB. However, the previously tested target genes for RNAi-mediated pest control (subsequently called *target genes*) exhibited moderate and slow-acting lethal effects. In this study, 27 double-stranded RNAs (dsRNAs) were orally delivered to identify highly effective target genes in CSFB adults by leveraging the findings of a genome-wide RNAi screen in *Tribolium castaneum*. Our screen using 500 ng of dsRNA identified 10 moderately effective (> 50% mortality) and 4 highly effective target genes (100% mortality in 8-13 days). The latter mainly included proteasome subunits. RT-qPCR experiments confirmed target gene silencing and dose-response studies revealed LD_50_ values as low as ∼20 ng in 14 days following a single exposure to dsRNA. Four highly effective dsRNAs also inhibited leaf damage (up to ∼75%) and one affected locomotion. The sequences of promising target genes were subjected to *in silico* target prediction in non-target organisms, e.g., beneficials such as honeybees, to design environmentally friendly dsRNAs. Overall, the study provides valuable insights for the development of dsRNA-based insecticides against CSFB.

## Introduction

The global human population is predicted to increase by ∼2 billion in the upcoming 3 decades, totaling up to 10 billion people (Gu et al., 2021). Hence, agricultural productivity must increase to meet the needs of the growing human population (van Dijk et al., 2021). A major hurdle in this regard is insect and other arthropod pests, whose damage may amount to ∼20% of the crop yields (Deutsch et al., 2018). Small-molecule insecticides, such as neonicotinoids, have been the pillar of insect pest control (Sparks et al., 2015). Although these insecticides have allowed effective control of most serious pests, they have also significantly contributed to biodiversity loss and an increase in detrimental chemical residues (Oosthoek, 2013). Moreover, numerous pest species have developed resistance to one or multiple insecticides, leading to reduced efficacy of crop protection (Feyereisen, 1995; Sparks & Nauen, 2015). These issues have prompted research into alternative pest management strategies, including RNA interference (RNAi).

The RNAi pathway is induced by double-stranded RNA (dsRNA) produced by viruses and transposons in nature. Upon the recognition of intracellular dsRNA by endonuclease Dicer-2, it is cleaved into ∼19 base pair RNA duplexes with 2 nucleotides 3’ overhangs, referred to as small interfering RNA (siRNA) (Kim et al., 2006; Liu et al., 2006; Naganuma et al., 2021). The siRNAs guide the cleavage of complementary mRNAs by the RNA-induced silencing complex (RISC), leading to gene-specific silencing (Martinez et al., 2002; Preall & Sontheimer, 2005). RNAi has emerged as a promising solution to insect pest control due to its species-specific and environmentally friendly characteristics (Baum et al., 2007; Mao et al., 2007; Liu et al., 2020). RNAi-based pest management depends on the delivery of dsRNAs that target genes involved in essential functions, such as proteasome-mediated proteolysis of proteins (Ulrich et al., 2015; Mehlhorn et al., 2021). Two prominent dsRNA delivery methods are either growing transgenic crops that transcribe dsRNA (e.g., SmartStax Pro maize against corn rootworms) (Reinders et al., 2023) or spraying externally produced dsRNA onto crops (e.g., Ledprona against Colorado potato beetle) (Rodrigues et al., 2021). The target pest should be susceptible to orally ingested dsRNA, and the most effective target genes in the target pest should be identified for either method to work effectively (Cedden & Bucher, 2024, this issue). Coleopterans and especially leaf beetles (Chyrsomelidae) generally show excellent responses to orally delivered dsRNAs (Bachman et al., 2013), making them prime targets for RNAi-based pest management (Baum et al., 2007; Bolognesi et al., 2012; Zhang et al., 2015; Willow & Veromann, 2021).

The cabbage stem flea beetle is an important pest of winter oilseed rape crops, particularly in northern Europe (Sivčev et al., 2016; Li et al., 2024). The larvae feed inside petioles and stems through tunneling, while the adults feed on the leaves and cotyledons of the crops (Ortega-Ramos et al., 2022). Managing CSFB has become increasingly challenging due to the ban on neonicotinoid seed treatments and the emergence of pyrethroid-resistant populations, the primary insecticide group used against CSFB in Europe (Højland et al., 2015; Willis et al., 2020; Ortega-Ramos et al., 2022; Li et al., 2024). In a previous study, we showed that RNAi has the potential to control CSFB (Cedden et al., 2024). The study showed that oral delivery of dsRNA leads to the generation of ∼21 nt siRNAs, systemic target gene silencing, and significant mortality. However, the highest mortality observed in the study was 76%, which was achieved within 25 days by dsSec23. Such performance under laboratory conditions might not suffice for inadequate pest control under field conditions, where limited by various environmental factors, such as UV light (Parker et al., 2019). In conclusion, more effective target genes had to be identified in CSFB for effective RNAi-based strategies as viable alternatives to pyrethroids.

Recently, a genome-wide RNAi screen has identified the most effective target genes for RNAi-mediated pest control in the red flour beetle *Tribolium castaneum* (Buer et al., 2024). About half of a subset of these genes were also effective when their orthologs were tested in the mustard beetle *Phaedon cochleariae* via oral dsRNA delivery. Furthermore, 11 out of the 12 effective target genes in the mustard beetle were also effective targets in the Colorado potato beetle *Leptinotarsa decemlineata*. These results suggest that the genes identified by Buer *et al*. (2024) are an excellent starting point for identifying effective target genes in other insects, especially coleopterans (Cedden & Bucher, 2024, this issue). In this study, we orally tested dsRNAs targeting the CSFB orthologs of 27 genes from the genes identified by Buer *et al*. (2024) to find very effective target genes against this pest. A subset of the highly effective dsRNAs was further investigated for dose-response relationships, effects on target gene expression, and behavioral effects (feeding and locomotion). Lastly, an *in silico* off-target prediction analysis was conducted to identify target regions suitable for environmentally friendly pest control.

## Results

### The oral RNAi screen identified effective target genes

In Buer et al. (2024), 34 superior target genes were identified to induce lethality in two beetle species, namely *T. castaneum* and *P. cochleariae*, by injection and oral application, respectively. Eleven of those had successfully been tested in *L. decemlineata*. To test for inter-specific transferability, we chose the CSFB orthologs of 14 genes from the 34 s*uperior target genes* from Buer et al. (2024) (indicated with stars in Fig. 1) and 13 from other effective target genes in *T. castaneum* (9 from cluster 1 and 4 from clusters 2-5, see supporting file 1, candidates) that were not previously transferred to pests in Buer et al. (2024). Most candidate target genes were categorized into three groups based on their cellular functions, namely proteasome, gene expression/translation, and secretion/endocytosis, whereas the rest were assigned into a miscellaneous category due to their diverse functions.

**Figure 1.**
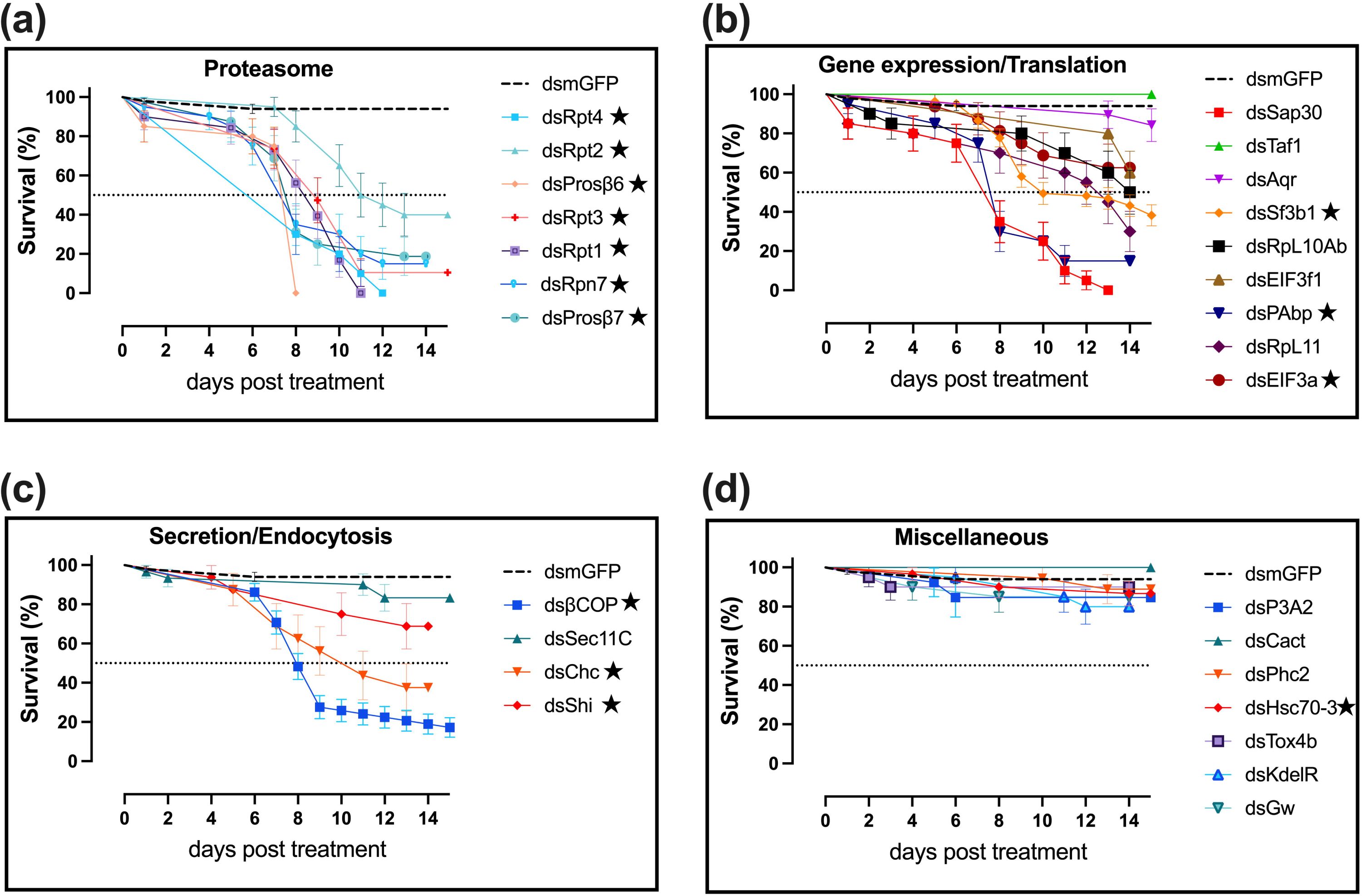
RNAi screen to identify efficacious target genes. Newly emerging adult cabbage stem flea beetles (sex-mixed) were initially fed with 500 ng of essential-gene targeting double-stranded RNAs (dsRNAs) or control dsRNA (dsmGFP), which were applied on top of oilseed rape leaf disks. Subsequently, the beetles were provided with untreated oilseed rape leaf disks ad libitum. The survivals of the beetles were observed daily for the following 2 weeks and investigated using Kaplan-Meier survival analysis. The curves show the survival probability percentages (%) ± standard error (the number of replications was 99 for the dsmGFP control treatment, and the number of replications for the other treatments ranged from 15-81, with an average of 23, see supporting file 2, survival 500 ng for details). The dsRNAs tested in the primary screen were grouped into 4 categories: proteasome (a), gene expression/translation (b), secretion/endocytosis (c), and miscellaneous (d). The survivals of dsmGFP-treated beetles (∼95% in all bioassays) throughout the primary screen were combined into a single survival curve and given as a reference in every panel (dashed black lines). The dsRNAs that were marked with a star (★) indicate that the target was present in the list of 34 target genes successfully transferred from a screen in *Tribolium castaneum* to the mustard beetle *Phaedon cochleariae* by Buer *et al*. (2024).

In the primary screen, candidate target genes were subjected to RNAi by feeding CSFB adults with a single dose of 500 ng of gene-specific dsRNA on oilseed rape leaf disks, which were consumed within one day. Subsequently, the survival of the adults was monitored daily for 14 days. Interestingly, all proteasome-related gene-targeting dsRNAs achieved at least a 50% reduction in survival within 14 days (Fig. 1a). Notably, dsProsβ6 was able to kill the whole tested population (n = 20) within 8 days (Fig. 1a), compared to maximum mortality of 76% after 25 days observed for the most effective treatment (200 ng dsSec23) in the previous study (Cedden et al., 2024). Interestingly, the median lethal time calculated for dsProsβ6 was also 8 days (Tab. S1), meaning that the population gave a homogenous response to the dsRNA. The other proteasome targeting dsRNAs that reached 100% mortality were dsRpt1 and dsRpt4, which had median lethal times of 9 and 8, respectively (Tab. S1). Although these median lethal times were comparable to that observed in dsProsβ6 treatment, 100% mortality was reached at later time points —specifically, 11 days post-treatment for dsRpt1 and 12 days post-treatment for dsRpt4.

Targeting genes related to gene expression/translation (including ribosomal proteins, Poly(A)-binding proteins, histone modifiers, etc.) has also identified some dsRNAs capable of above 50% mortality. Only one of the dsRNAs in this category, namely dsSap30, was able to achieve 100% mortality, which took 13 days with a median lethal time of 8 days (Fig. 1b and Tab. S1). Other dsRNAs in this category that caused a notable reduction in survival were dsPAbp, dsSf3b1, dsRpL11, and dsRpL1Ab, which caused 85%, 73.2%, 70%, and 50% mortalities, respectively. Two targets in the secretion/endocytosis category were able to reduce the survival of the beetles below 50%. dsβCOP and dsChc, achieved 82.8% and 62.5% mortalities in 14 days (Fig. 1c and Tab. S1). None of the targets in the miscellaneous category caused a notable reduction in survival (Fig. 1d).

The hazard ratios (dsmGFP/essential-gene targeting dsRNA) of dsRNAs causing 100% mortality were calculated to be ∼20, with the dsSap30 having the highest ratio of 24.3 (Tab. S1). As expected, the survival curves of the effective dsRNA treatment groups significantly differed from that of the dsmGFP group (P < 0.003, Tab. S1). Overall, the primary screen showed that 14 out of 27 tested dsRNAs achieved at least 50% mortality and 4 of them, namely dsRpt4, dsProsβ6, dsRpt1, and dsSap30, achieved 100% mortality within 14 days at 500 ng dsRNA. Of these top target genes, all but *sap30* were included in the 34 superior target genes from Buer et al. (2024). Together with the results of the other tested genes, this confirms the previous observation that any given target gene can have different efficacies in different species (Cedden & Bucher, 2024, this issue, Mehlhorn et al., 2021).

### Dose-responses to effective dsRNAs

The lethal effects of a subset of effective dsRNAs dsRpt1, dsRpt4, dsSap30, dsProsβ6, and dsProsβ7 were further tested at lower doses to investigate dose-response relationships (Fig. 2). Initially, we tested these five dsRNAs at 50 ng, i.e. a 10-fold dilution. The results showed that only dsRpt1, dsRpt4, and dsProsβ7 could lower the survival of the beetles below 50% within 14 days (Fig. 2a). The median times of survival for these three treatments were 8.5, 8, and 12 days post-treatment, respectively. Hence, we further tested these three dsRNAs at 5 ng to estimate their LD_50_ (median lethal dose) values. The logistic regression analysis showed that the LD_50_ values for dsRpt1, dsRpt4, and dsProsβ7 were 20.7 ng (95% CI: 9.2-31.7), 18.87 ng (95% CI: 9.9 to 36), and 232.1 ng (95% CI: 136.9 to 375.0), respectively, when the endpoint was 14 days post-treatment mortality (Fig. 2b). The areas under the receiver operating characteristic curves (ROC) for the logistic regression analyses were 0.91, 0.93, and 0.87, respectively, suggesting that the models had good predictive capabilities (Fig. S1). The study results indicated that CSFB exhibited variable dose-responses to different dsRNAs. Notably, two proteasome-targeting dsRNAs, dsRpt1 and dsRpt4, had the lowest LD_50_ of ∼20 ng.

**Figure 2.**
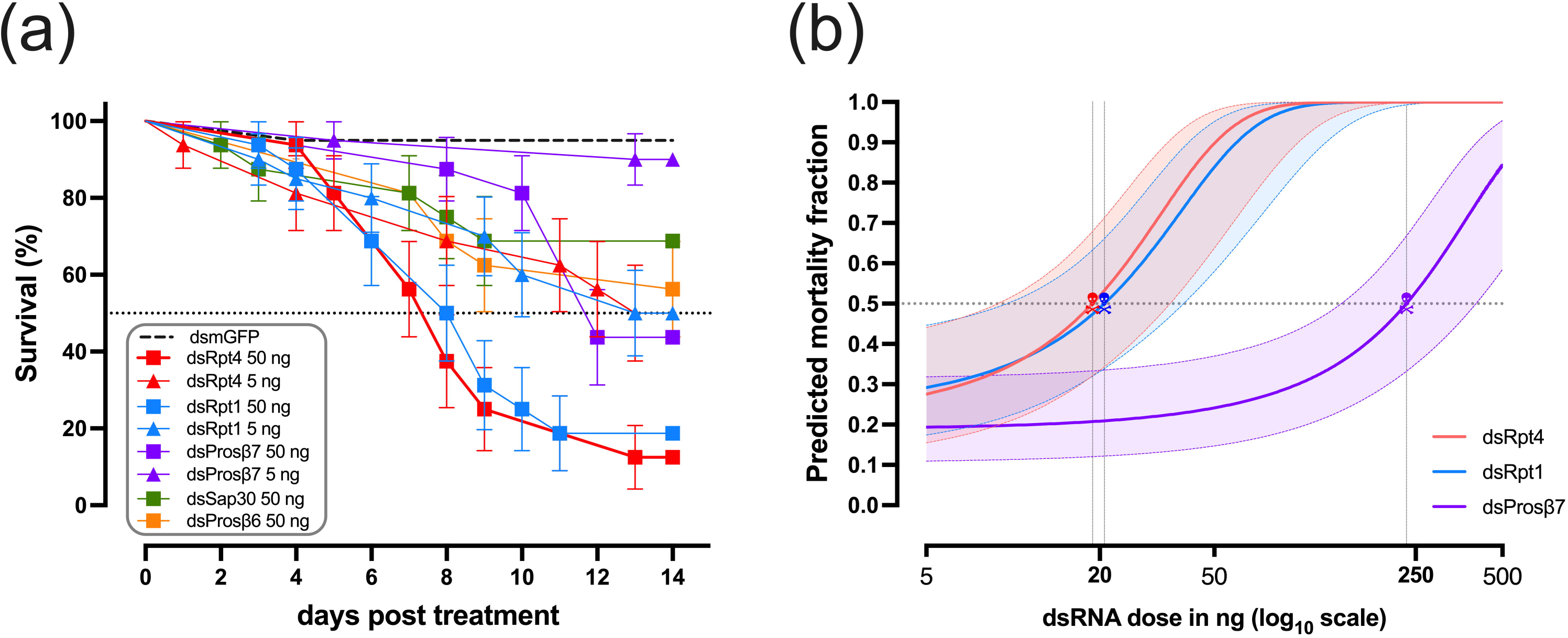
Dose-responses to dsRNAs treatments. (a) Five dsRNAs that were highly lethal in the primary screen at 500 ng were selected for testing at 50 ng. The three dsRNAs that caused at least 50% mortality in 14 days post-treatment at 50 ng were tested at a further reduced dose of 5 ng. The curves show the survival probability percentages (%) ± standard error (n= 16-20). (b) Three dsRNAs were subjected to logistic regression analysis by inputting the total mortality in 14 days post-treatment at 500 ng, 50 ng, 5 ng, and 0 ng (mortality in the dsmGFP group) to estimate the LD_50_ (median lethal dose) values (indicated with skulls). The plot shows the predicted mortality fractions at different dsRNA doses ± 95% confidence interval.

### Gene silencing induced by effective dsRNAs

An RT-qPCR experiment was conducted to investigate to which degree the expressions of the effective target genes were affected by the oral delivery of 500 ng dsRNA at 3 days post-treatment (i.e., putatively during the mounting of an RNAi response but before observation of phenotypes). All 4 tested dsRNAs achieved significant silencing of their respective targets in the CSFB whole bodies (Fig. 3). Interestingly, three proteasome targeting dsRNAs, namely dsRpt1 (73%, P < 0.001), dsRpt4 (49%, P = 0.014), and dsProsβ6 (54%, P = 0.01) achieved more silencing compared to the dsSap30 (38%, P = 0.028), all relative to the expression in the dsmGFP control. Hence, target gene-specific variability in % silencing efficiency was present. The variability in gene silencing was partially correlated with the basal expression (data from RNA-seq in intact CSFB females) of the respective genes, where higher basal expression was associated with better silencing efficiency in % (Fig. S3). There was a 94.9% (P = .026) positive correlation between the % silencing efficiencies and the basal expressions, albeit 4 data points are not enough for a firm conclusion. For instance, dsRpt1 caused the highest percent silencing, and the mean read count of its target was 3818 TPM (median transcripts per million, n = 7), while dsSap30 caused the lowest percent silencing and the mean read count of its target gene was 471 TPM (Fig. 3). Notably, strong lethal effects observed in the primary screen do not seem to depend on high silencing efficiencies. For instance, 500 ng dsSap30 caused merely 38% gene silencing, while the same dose led to 100% mortality in 13 days (Fig. 1b) with the highest hazard ratio (ratio: 24.3, Tab. S1). The results show that effective dsRNAs caused silencing with variability that is positively correlated with basal target gene expression, although the degree of silencing does not seem to be relevant for lethality.

**Figure 3.**
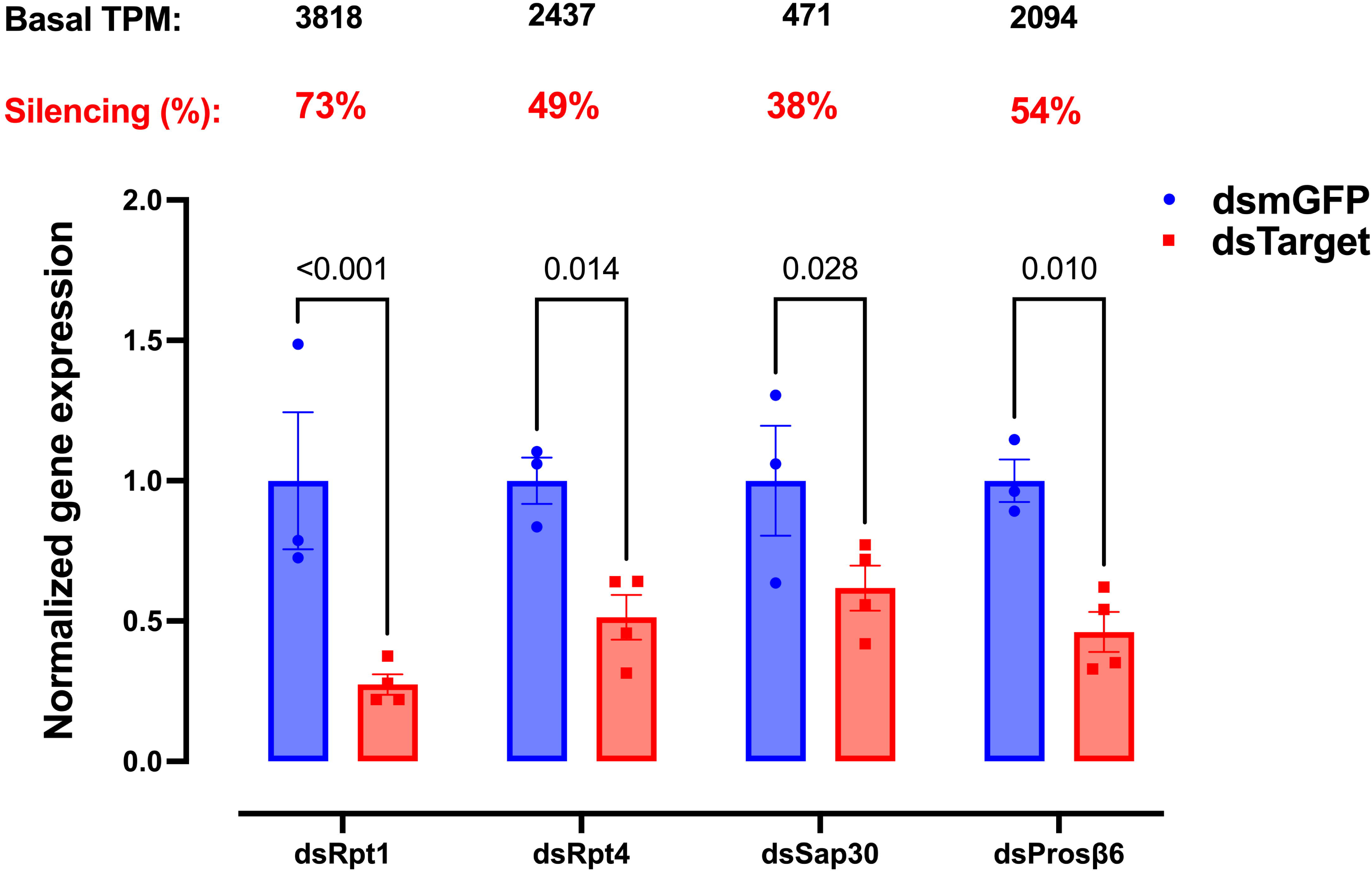
Reduction of gene expressions after dsRNA treatments. Each dsRNA was orally delivered at 500 ng to newly emerged CSFB adults, which were sampled 3 days post-treatment. The bars show the mean ± SEM (n = 3 – 4, each replication contained homogenized whole bodies of one male and one female) normalized expressions of the genes targeted by the respective dsRNA. The expression in the dsmGFP treatment group was used to normalize the target gene expression. The expression of the target gene in the respective dsRNA treatment group was compared to the expression in the dsmGFP control group using multiple t-tests corrected for multiple comparisons using the Holm-Šídák method. The percentages above the bars show the percent reductions in the expression of the target gene in the respective dsRNA treatment group. The basal transcripts per million (TPM) values indicate the median expressions of the respective genes in intact CSFB females measured through RNA-seq (n = 7, data is available at https://doi.org/10.6084/m9.figshare.24085815).

### Behavioral effects of lethal dsRNAs

Previously, we showed that essential-gene targeting dsRNAs can have various sublethal effects on CSFB (Cedden *et al*., 2024). Here, we investigated whether the effective dsRNAs affect the locomotion and feeding behaviors of the adult CSFB (Fig. 4). Two of the most effective dsRNAs in the primary screen (dsProsβ6: fastest 100% lethality in the primary screen and dsRpt4: lowest LD_50_) were selected for 2 h movement tracking at 4 days post-treatment. The results showed that dsRpt4 significantly reduced the mean total movement by 74.7% (P = .002, Z = 3.30, n = 6) (Fig. 4a). Although dsProsβ6 had reduced the mean total movement by 59.3%, this effect was not significant (P = .117, Z = 1.89, n = 6). The beetles’ walking speeds did not differ significantly between the dsRNA treatment groups (P > 0.05) (Fig. 4b). Hence, the propensity to walk rather than the actual movement capacity of the beetles seems to be affected by the dsRpt4 treatment as only the total movement, not the speed, was reduced.

**Figure 4.**
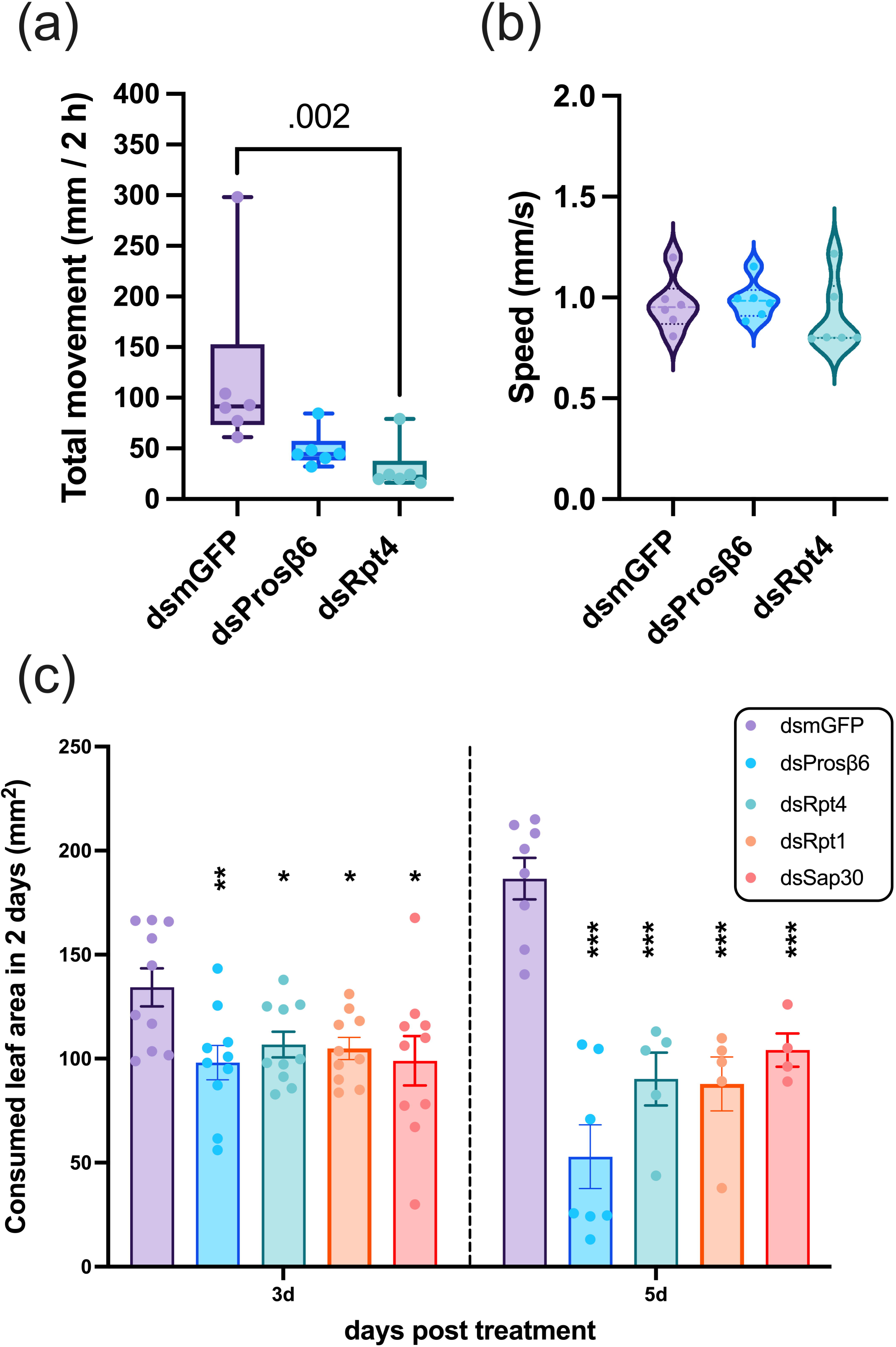
Behavioral effects of dsRNA treatments. Each beetle orally received 500 ng of dsRNA treatment upon emergence. (a) Total movement (mm per 2 hours) of beetles (median, quartiles, and range are shown in box plots, n = 6) in 2 hours was investigated using the Zantiks LT system 4 days post-treatment. The Kurskal-Wallis test, followed by Dunn’s tests, was performed to compare the total movement in two essential-gene targeting dsRNA treatment groups with that of the dsmGFP control group; adjusted P values < 0.05 are shown (see supporting file 2, total movement). (b) The speed (mm/s) of beetles (violin plots, n = 6) derived from the same movement-tracking data by dividing non-zero real-time movement data by 5 seconds (bin value of our tracking set-up). The Kurskal-Wallis test yielded a P value of 0.4, and no pair-wise tests were conducted (see supporting file 2, speed). (c) The consumed leaf area by beetles (mean ± SEM, n = 5-10) at 3 and 5 days post-treatment, reflecting the consumption in the preceding 2-day period. Multiple t-tests with the Holm-Šídák corrections were performed. The level of significance of the difference between the essential-gene targeting dsRNA group and the dsmGFP control group was indicated above the bars (*P < 0.05, **P < 0.01, ***P < 0.001) (see supporting file 2, leaf consumption).

The leaf consumption was investigated 3 and 5 days post-treatment, reflecting the consumptions in the preceding 2-day periods. The leaf consumption was significantly affected at both 3 days and 5 days post-treatment by all the tested dsRNAs (dsProsβ6, dsRpt4, dsRpt1, and dsSap30) compared to the dsmGFP control (P < .05, Fig. 4c). This effect increased over time in all dsRNA treatments with a rather mild reduction 1-3 days post-treatment (27.0%, 20.5%, 21.9%, and 26.3% reductions, respectively) but with a strong reduction 3-5 days post-treatment (71.7%, 52.9%, 51.6%, and 44.2% reductions, respectively). Of note, the increase in the feeding of dsmGFP-treated beetles observed from 3 to 5 days (38.9% increase) reflected the usual feeding behavior of CSFB after emergence as the feeding rate spontaneously peaks in ∼6-day-old CSFB adults (Güney et al., in preparation). The feeding inhibition effects of the tested dsRNAs were very similar at 3 days post-treatment, but some variability was observed at 5 days post-treatment. dsProsβ6 treatment achieved the highest mean feeding inhibition of 72% (P < .001, df = 13) at 5 days post-treatment. The results indicate that effective dsRNAs are able to reduce leaf damage even before lethality occurs.

### Off-target analysis of effective target genes

Our primary screen and subsequent experiments revealed that dsProsβ6 (100% mortality and high feeding inhibition), dsRpt4 (low LD_50_ and reduced locomotion), dsRpt1 (low LD_50_), dsSap30 (100% mortality), and dsProsβ7 (moderate LD_50_) are promising dsRNAs for CSFB management. As a second task, we aimed to identify target gene sequences that were safest in non-target organisms (i.e., beneficial) based on *in silico* off-target prediction in key environmental invertebrate species. To that end, we selected western honey bee *Apis mellifera*, buff-tailed bumblebee *Bombus terrestris*, seven-spot ladybird *Coccinella septempunctata*, *Daphnia magna*, and common green lacewing *Chrysoperla carnea*. This selection included species that interact with the oilseed rape crops and species that are usually tested during the registration of insecticides.

The analysis identified many hits in the transcriptomes of non-target organisms with gene-dependent variability (left panels in Fig. 5). Interestingly, 3 target genes had off-target hotspots with more than 50 gene hits in multiple off-target species. Such hotspots were not observed in *Pchr-prosβ6* and *Pchr-prosβ7* (left panels in Fig. 5a,e). All these hotspots included hits to *A. mellifera*, an important pollinator. Subsequently, we focused on hits to genes that were orthologous to the essential genes (>90% mortality upon injection in *T. castaneum*) identified by Buer *et al*. (2024). The number of target gene ortholog hits ranged from 0 to 7 (right panels in Fig. 5). *Pchr-prosβ6* and *Pchr-prosβ7* genes had the lowest number of overall and per siRNA essential gene hits. Moreover, they had > 100 bp regions without any target gene hits that would be optimal regions to target in terms of minimizing the detrimental effects on off-target organisms. The complete and detailed list of hits is available on FigShare (https://doi.org/10.6084/m9.figshare.25283683). Based on this analysis, we designed 450 bp target sequences with minimal essential gene off-targets in the analyzed non-target species (provided in supporting file 1, off-target reduction). The new dsRNAs designed to target *Pchr-prosβ6, Pchr-rpt1, Pchr-sap30*, and *Pchr-prosβ7* had less than 50 essential gene hits (47, 43, 19, and 24, respectively). However, the new dsRNA designed to target *Pchr-rpt4* still contained many essential gene off-targets (196 targets). The results showed different gene sequences have varying potentials for designing off-target minimized dsRNAs, suggesting another criterion for determining good RNAi target genes beyond lethal and sublethal effects in the target pest.

**Figure 5.**
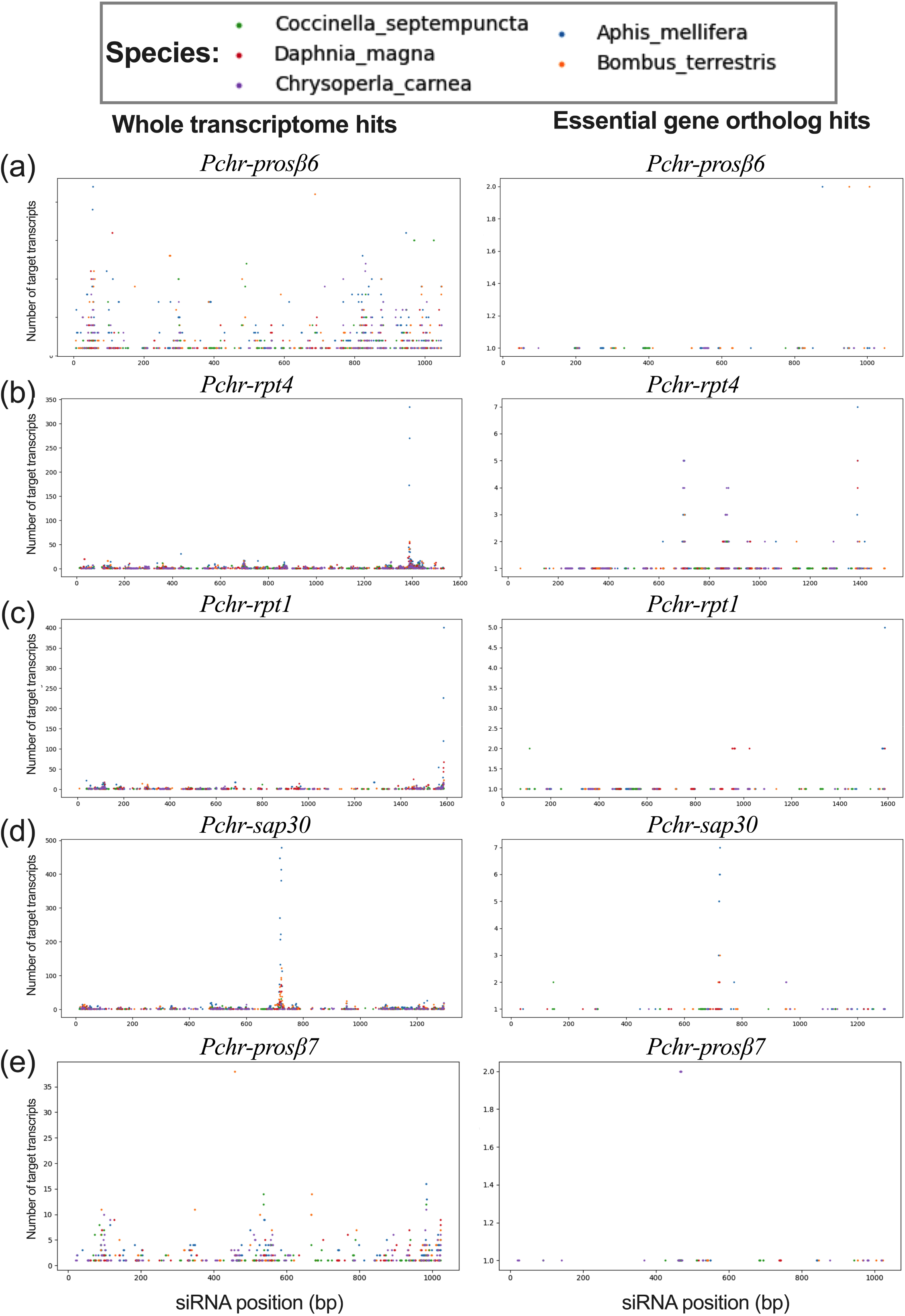
*In silico* prediction of off-targets in other organisms. The mRNA sequences of the top-performing 5 target genes identified in the cabbage stem flea beetle adults, namely *Pchr-prosβ6* (a), *Pchr-rpt4* (b), *Pchr-rpt1* (d), *Pchr-sap30* (d), *Pchr-prosβ7* (e) (see supporting file 1, gene accessions) were converted into 21-mer small interfering RNA (siRNA) sequences and mapped onto the transcriptomes of non-target species *Apis mellifera, Bombus terrestris, Coccinella semptepunctata, Dapnia magna, Chrysoperla carnea* using Bowtie 1 v1.3. The X-axis shows the location of the first nucleotide of the siRNA on the respective gene sequence in the cabbage stem flea beetle, and the Y-axis shows the number of gene matches (hits) per siRNA. The color of the dots represents the non-target species. The left panels show hits to any gene in the transcriptomes of the non-target species. The right panels show the same analysis restricted to ∼900 orthologs of genes that were essential in the red flour beetle (above 90% lethality in *Tribolium castaneum* genome-wide screen, Buer *et al*. (2024). Based on this analysis, we designed target sequences of 450 bp length with reduced number of essential gene off-targets in these species (provided in supporting file 1, off-target reduction).

## Discussion

Identification of effective target genes is a key step in the development of RNAi-based pest management strategies (Mehlhorn et al., 2021). In this work, we aimed to identify highly effective target genes by testing the CSFB orthologs of the effective target genes established by Buer *et al*. (2024) based on a genome-wide RNAi screen in *Tribolium castaneum*. Indeed, this approach proved successful as the target genes identified here had more promising lethal effects compared to those investigated previously by Cedden *et al*. (2024). For instance, dsRpt4 had an LD_50_ of 18.8 ng in this work, whereas dsSec23 had an LD_50_ of 87 ng (Cedden et al., 2024), even though the endpoint was 10 days longer in the latter study.

### Selecting candidate target genes for testing

Approximately half of the candidate target genes tested in this study (14/27) were part of a list of *superior target genes* first identified in the red flour beetle *T. castaneum* and subsequently confirmed to cause high lethality when tested through oral dsRNA delivery in the mustard beetle *Phaedon cochleariae* (Buer *et al*., 2024). We observed that 79% (11/14, Tab. S1) of the effective target genes identified in CSFB overlapped with these genes confirmed in *P. cochleariae*. The high success rate highlights the usefulness of the *superior target gene* list provided by Buer *et al*. (2024) when searching for effective target genes in other insect species. A notable exception was the results obtained with dsHsc70-3. While *hsc70-3* was an effective target in both *P. cochleariae* and *L. decemlineata* (Buer *et al*., 2024), dsHsc70-3 did not have any observable effect on the survival of CSFB. On the other hand, we identified 3 effective targets, namely 60S ribosomal L10a, 60S ribosomal L11, and SAP30-binding, that were not previously transferred to other pests by Buer *et al*. (2024), albeit other ribosomal proteins were included in the *superior target genes*. The discrepancies might be explained by the differences in the stages used during the bioassays (adults of CSFB versus larvae of other species) and the existence of functional redundancy for the gene, depending on the species (Cheatle-Jarvela et al., 2020; Cedden & Bucher, 2024, this issue).

### The proteasome is an excellent target for RNAi-based pest control

Our oral RNAi screen suggested that proteasome-related gene targets might be especially ideal targets, at least in CSFB adults. All tested proteosome-related targets (7 genes) achieved at least 50% mortality, suggesting that testing further proteosome-related genes might also be beneficial. Interestingly, the first phase of the genome-wide RNAi screen in *T. castaneum* also identified proteasome as the prime target in *T. castaneum* (Ulrich et al., 2015). In addition, the first dsRNA-based sprayable insecticide, namely Ledprona, that is currently entering the market is also targeting a proteasome subunit (Rodrigues et al., 2021). Hence, targeting the proteasome is becoming understood as an effective RNAi-based mode of action against insects and our results strengthen this notion. However, given that we tested only a subset of *superior target genes* and selected the proteasome for that purpose, it cannot be excluded that parts of the prominence of the proteasome as a target is due to a selection bias. Other pathways or genes may prove to have comparable or even superior effects in CSFB or other species.

Interestingly, no commercial small-molecule insecticides currently target the proteasome (Sparks & Nauen, 2015; Sparks et al., 2020). It could be that depleting the mRNAs of proteasome subunits is superior to inhibiting their protein products – the mode of action of conventional insecticides. Counterintuitively, the proteasome is known to be a stable complex, at least in the rat liver, with a half-life of up to 15 days (Tanaka & Ichihara, 1989; Pack et al., 2014; Marshall & Vierstra, 2019). This is in contrast to the general assumption that the protein of a good RNAi target should be unstable for it to be sufficiently susceptible to mRNA depletion (Knorr et al., 2021). Hence, other factors such as mRNA turnover and the proteasome’s vital function might be making the proteasome a very effective RNAi target. Alternatively, small molecule inhibitors of proteasome might have adverse effects on non-target organisms and might not pass safety controls. Targeting the proteasome at the protein level through the application of conventional insecticides might not be feasible due to the highly conserved structure of the complex, increasing detrimental effects on non-target organisms. In contrast, our study revealed that effective proteasome targets such as *Pchr-prosβ6* and *Pchr-prosβ7* contained many regions lacking off-targets in non-target organisms. This suggests that designing bio-safe and eco-friendly insecticidal dsRNAs targeting proteasomes is possible.

### Are genes with higher basal expression better targets?

A previous study systematically studied the link between the basal mRNA expression and silencing efficiency of RNAi in three species and found a positive correlation (Chen et al., 2021). In line with this observation, our results also suggest a positive correlation as genes with higher basal expressions were more effectively silenced in CSFB whole bodies. Importantly, however, it is unclear whether silencing efficiency, which is almost exclusively inferred through RT-qPCR measurements, is an insightful criterion for determining effective RNAi target genes for pest management (Mehlhorn et al., 2021). In the current study, there was a high variability between the silencing efficiencies (38% to 73%) of the most effective dsRNAs, albeit all still caused 100% mortalities at 500 ng. The silencing efficiencies might be misleading because RT-qPCR also measures pre-mRNA (Zeisel et al., 2011), which might be upregulated after RNAi by regulatory gene networks that try to rescue the suppressed transcript levels (Mehlhorn et al., 2021), and the threshold required to induce lethality might be gene-specific rather than a general threshold, making comparisons between different target genes challenging. Given the variability between species and further uncertainties, it seems the best strategy to test several candidate target genes by oral dsRNA delivery instead of putting too much effort into predicting efficacy (see Cedden & Bucher, 2024, this issue, for more extensive discussion).

### Targets to be used for application in the cabbage stem flea beetle

RNAi screens are capable of identifying many effective targets in a pest, as demonstrated in the current and in earlier studies (Ulrich et al., 2015; Knorr et al., 2018; Buer et al., 2024). However, further steps in the pesticide development pipeline, such as field trials and safety assessment, are restricted to a limited number of active ingredients (e.g., dsRNA) due to economic considerations. That is why the investigation of all relevant behavioral effects and *in silico* off-target analysis of promising target genes are required to select gene targets for further development instead of a sole focus on lethal effects. For instance, dsProsβ6 had up to 75% feeding inhibition effect within 5 days post-treatment. Hence, crop protection might start days before the lethality of the animals kicks in after approximately seven days. A similar observation was made by Mehlhorn et al., (2021) in the *P. cochleariae* orally treated with a dsRNA targeting the proteasome. The observation that dsRNA treatments lead to feeding inhibition before lethality indicates that in field applications, both parameters will influence overall pest control efficacy.

On the other hand, the most promising dose responses were observed in the bioassays with dsRpt1 and dsRpt4, which had LD_50_ values of ∼20 ng, corresponding roughly to 7 g/ha. Taking all our data together, we suggest that the two proteasome subunit targets, Pchr-*rpt1* and *Pchr-rpt4*, might be considered for further developing efficient spray-based dsRNA insecticides against CSFB. These targets had behavioral effects in addition to excellent dose-responses, suggesting they may provide the highest crop protection in the field. However, the off-target hotspots revealed by our *in silico* analysis for these targets should be avoided to minimize potential detrimental effects on the non-target organisms. Based on this consideration, we designed target sequences (provided in supporting file 1, off-target reduction) that may be used in future RNAi-based CSFB management efforts.

## Experimental Procedure

### Insects

Cabbage stem flea beetle (CSFB, *Psylliodes chrysocephala*) laboratory colony was reared at 21° ± 1° C and 65 ± 10 % relative humidity under 16:8 light: dark cycle on winter oilseed rape plants (growth stage: BBCH 30-35) in rearing chambers. Freshly eclosed adults emerging from the soil were collected daily for bioassays or placed into the rearing chambers to maintain the colony. Field-collected larvae (Göttingen, Germany, experimental oilseed rape field without insecticide application, coordinates: 51.564065, 9.948447) were annually introduced into our laboratory colony to maintain genetic diversity.

### Selection of target gene candidates

We used our *de novo* transcriptome assembly representing various adult stages of female CSFB with a BUSCO v5.4.2 (Manni et al., 2021) completeness score of 95.9% (S:94.5%, D: 1.4%, F: 1.8%, M: 2.3%, n: 2124, comparison against the endopterygota v4 dataset) (Güney *et al*., 2024, NCBI BioProject: PRJNA930726, TSA: <u>GKIH01000001:GKIH01098921</u>) to identify candidate gene sequences. Orthofinder v2.5 (Emms & Kelly, 2019) with the default parameters was used to conduct orthology inference between the translated amino acid sequences from CSFB and *Tribolium castaneum* (Tcas5.2 from www.ibeetle-base.uni-goettingen.de). The longest amino acid sequences from the transcriptomes were extracted using TransDecoder v5.6 (github.com/TransDecoder) for the analysis. The CSFB orthologs of the target genes from Buer et al. (2024) were filtered, and corresponding transcript sequences of 27 candidate target genes were retrieved from the CSFB transcriptome (see supporting file 1 for NCBI accession numbers).

### dsRNA synthesis

The lengths of dsRNAs were within the range of 315 to 453 bp based on the lengths of the amplicon regions targeted by the gene-specific primers (dsRNA sequences were provided in supporting file 1, dsRNA primers). The dsRNAs were named after the gene symbols of *T. castaneum* orthologs (available at www.ibeetle-base.uni-goettingen.de) of the respective target genes in CSFB. Additionally, a control dsRNA named dsmGFP was synthesized using a DNA template synthesized by IDT (Germany). The candidate gene-specific dsRNAs were designed to target the coding sequence regions without predicted off-targets to other candidate target genes in CSFB (all CSFB orthologs of good target genes in Buer *et al*., 2024, see supporting file 1, orthology). The region was skipped if there was at least a 19 bp match with another target gene ortholog. In cases where the allowed target region became shorter than 315 bp, 5’ or 3’ UTR regions were also included in the target region.

Primer3 with the default parameters was used to design a pair of gene-specific primers to amplify the regions to be targeted by the dsRNAs, and the T7 promoter “GAATTGTAATACGACTCACTATAGG” was added to the 5’ ends of the primer pair (see supporting file 1, dsRNA primers). The primer pairs with and without the T7 promoter sequence were provided by Integrated DNA Technologies (IDT, Germany). The cDNA to be used as a template for PCR amplification was obtained from a mix of ten 1 to 25 days-old CSFB adults by RNA extraction using Quick tissue kit (Zymo) and reverse transcription using the LunaScript® RT SuperMix Kit, according to the manufacturer’s instructions. Two PCR reactions were performed per dsRNA; for the sense strand, T7 + forward and reverse primers were mixed, and for the antisense strand, the forward and T7 + reverse primers were mixed (0.5 µM). The 50 µL PCR reactions were performed using Q5® Hot Start High-Fidelity 2X Master Mix according to the manufacturer’s instructions (the annealing temperature was 56-60 °C, and 28-30 cycles were performed). Single bands with the expected length were extracted following agarose gel electrophoresis using the Gel and PCR Clean-up kit (MachereyLJNagel, Germany) to be used for the transcription reactions. The sense and antisense RNA strands were synthesized separately in 20 µL reactions using MEGAscript™ T7 Transcription Kit (Invitrogen) and purified through Lithium Chloride precipitation. Two RNA strands were equimolecularly combined in nuclease-free water by measuring the concentrations using a nanodrop (40 μg/OD260 conversion factor). The RNA strands were annealed by denaturation at 94 °C for 5 min, followed by cooling at room temperature for 30 min. The annealed dsRNAs (2.5 µg) were checked on 1.5% agarose gel by running them together with 2 µL of dsRNA ladder (NEB# N0363S, Germany) (Fig. S2).

### Bioassays

The dsRNAs were diluted to their test doses per µL (e.g., 500 ng/µL for the primary screen) in nuclease-free water and combined with Triton-X (200 ppm, Sigma-Aldrich, Germany). Each Petri dish (60 x 15 mm with vents) was prepared as an independent replication and included one newly emerged adult CSFB (sex-mixed) and 30 mm^2^ leaf disk punctured from the first true leaves of oilseed rape plants (growth stage: BBCH ∼35) placed on top of 80 mm^3^ 1% agarose gel. One µL from the dsRNA solution was pipetted and homogenously spread across the surface of the leaf disk, which was allowed to dry for 10 min before adding the beetles. The Petri dishes were kept in the rearing chambers described above. After 24 h, the old agarose gel disks were replaced with untreated 400 mm^2^ leaf disks placed on top of agarose gel disks, and this replacement was repeated every third day for the following 14 days. The survival of the individual beetles was checked daily, as described by Cedden *et al*. (2024). The number of replications was 99 for the dsmGFP control treatment, and the number of replications for the other treatments ranged from 15-81, with an average of 23 (see supporting file 2, survival 500 ng for the number of replications and raw survival data).

### Gene expression measurement

The newly emerged CSFB (n = 3-4, each replication consisted of homogenized whole bodies of one male and one female) receiving 500 ng dsRNA per leaf disk treatment (as described in the Bioassays section) were flash frozen using liquid nitrogen and stored at -80 °C immediately after removing their wings. The total RNA from CSFB samples was extracted using the Quick-RNA Tissue & Insect MicroPrep™ Kit (Germany) according to the manufacturer’s instructions. RT-qPCR was performed using the Luna One-step RT-qPCR kit (New England Biolabs, Germany) on a 384 well-plate with 3 technical replicates per biological replicate in CFX384 Touch Real-Time PCR (Bio-Rad) according to the manufacturer’s instructions. Each well contained 0.8 µL of the total RNA (500 ng / µL) and 0.8 µL of 10 mM mix of forward and reverse primers (provided by IDT) to measure the expressions of dsRNA target genes (efficiencies of the primer pairs were 94-102%) and a reference gene, namely, *rps4e*, which was previously optimized for RNAi experiments (Cedden *et al*., 2024). The primers were designed using Primer3 to target regions that do not overlap with the respective dsRNA target sequence (see supporting file 1, qPCR primers). ΔΔCq values normalized to target gene expression in dsmGFP-fed samples were used for further analysis.

### Leaf consumption and movement behavior

The newly emerged CSFB (n = 10, sex-mixed adults) receiving 500 ng essential-gene targeting dsRNA or dsmGFP treatment were provided with 400 mm^2^ untreated leaf disks first after the consumption of the dsRNA-treated leaf disk (1 day post-treatment) and then 2 days later following the removal of the previous leaf disk. The analysis detailed in Cedden et al. (2024) was used to calculate the leaf area consumptions during 1-3 and 3-5 days post-treatment. The beetles that died before a time point were excluded from the analysis.

The newly emerged CSFB (n = 6) receiving 500 ng essential-gene targeting dsRNA or dsmGFP treatment were kept as described in the bioassays section until 4 days post-treatment. Subsequently, the beetles were individually transferred to larger empty Petri dishes (100 mm x 15 mm with vents) for movement tracking using a Zantiks LT system (Cambridge, United Kingdom) with a 4×5 layout. The script used for the measurement with Zantiks LT is available on FigShare (https://doi.org/10.6084/m9.figshare.24162525.v1). Briefly, each beetle was independently tracked for 2 hours with the upper white lights on and in room temperature conditions with green under lights to stimulate foraging. The real-time movement was obtained with bin times of 5 seconds, which were summed per CSFB to get total movement in mm. The non-zero movement bins were divided by 5 seconds and averaged per individual beetle to obtain speed (mm/s) values per beetle.

### Off-target analysis

Off-target analysis was conducted using Bowtie v1.3 (Langmead et al., 2009) to map every possible 21-mers (i.e., potential siRNAs) of effective target gene sequences in CSFB. Mappings in both directions were allowed with up to 2 mismatches. Furthermore, the orthology inference analysis described above was conducted using translated amino acid sequences of *T.castaneum* and the transcriptomes of the investigated non-target organisms *Aphis mellifera* (RefSeq: GCF_003254395.2), *Bombus terrestris* (RefSeq: GCF_910591885.1), *Daphnia magna* (RefSeq: GCF_020631705.1), *Coccinella septempunctata* (RefSeq: GCF_907165205.1), and *Chrysoperla carnea* (RefSeq: GCF_905475395.1). The orthology inference results were filtered to get the orthologs of the target genes associated with above 90% mortality in Buer et al. (2024). The filtered list and the off-target prediction results were investigated to determine the number of matches to the putative essential gene orthologs in the non-target organisms. The Python 3.9 script used during data analysis and the detailed results are available at FigShare (https://doi.org/10.6084/m9.figshare.25283683).

### Statistical analysis

The Kaplan-Meier survival curves were plotted using the bioassay results. The curves of dsRNA treatment groups in the primary screen that reached 50% mortality in 14 days were compared with that of dsmGFP group using pairwise log-rank tests with Bonferroni correction. The dose responses were analyzed by fitting the mortalities in 14 days at different dsRNA doses (5 ng, 50 ng, and 500 ng for effective dsRNAs, while bioassay result with 500 ng dsmGFP treatment was used as 0 ng) into logistic regression models. The Kurskal-Wallis test, followed by Dunn’s tests, was performed to compare the total movement and speed data generated by the Zantiks LD behavioral analysis. Multiple t-tests with the Holm-Šídák corrections were performed to compare leaf consumption and gene expression data. Results were considered statistically significant at P < 0.05. All statistical procedures were conducted using GraphPad Prism v10.1.

## AUTHOR CONTRIBUTION

**Doga Cedden:** Conceptualization; investigation; writing – original draft; writing – review and editing; visualization; methodology; software; validation; formal analysis; project administration; data curation; supervision. **Gözde Güney:** Investigation; validation; data curation; formal analysis. **Xavier Debaisieux:** Investigation; validation. **Stefan Scholten:** Funding acquisition; methodology; resources. **Michael Rostás:** Funding acquisition; methodology; resources; supervision. **Gregor Bucher:** Conceptualization; methodology; writing – review and editing; supervision; resources; project administration.

## DATA AVAILABILITY STATEMENT

The data are available in the supporting information and on FigShare (https://doi.org/10.6084/m9.figshare.25599138.v1). Corresponding authors can be contacted for further details.

## ACKNOWLEDGEMENTS

Doga Cedden and Gözde Güney were funded by the Deutscher Akademischer Austauschdienst research grants program during the project. Open Access funding enabled and organized by Projekt DEAL. The authors thank Dr. Bernd Ulber, Dr. Gerd Vorbrüggen, Claudia Hinners, Jonas Watterott, and Derya Balci for their help in the project. The graphical abstract was created with BioRender.com with support from Metin Cedden.

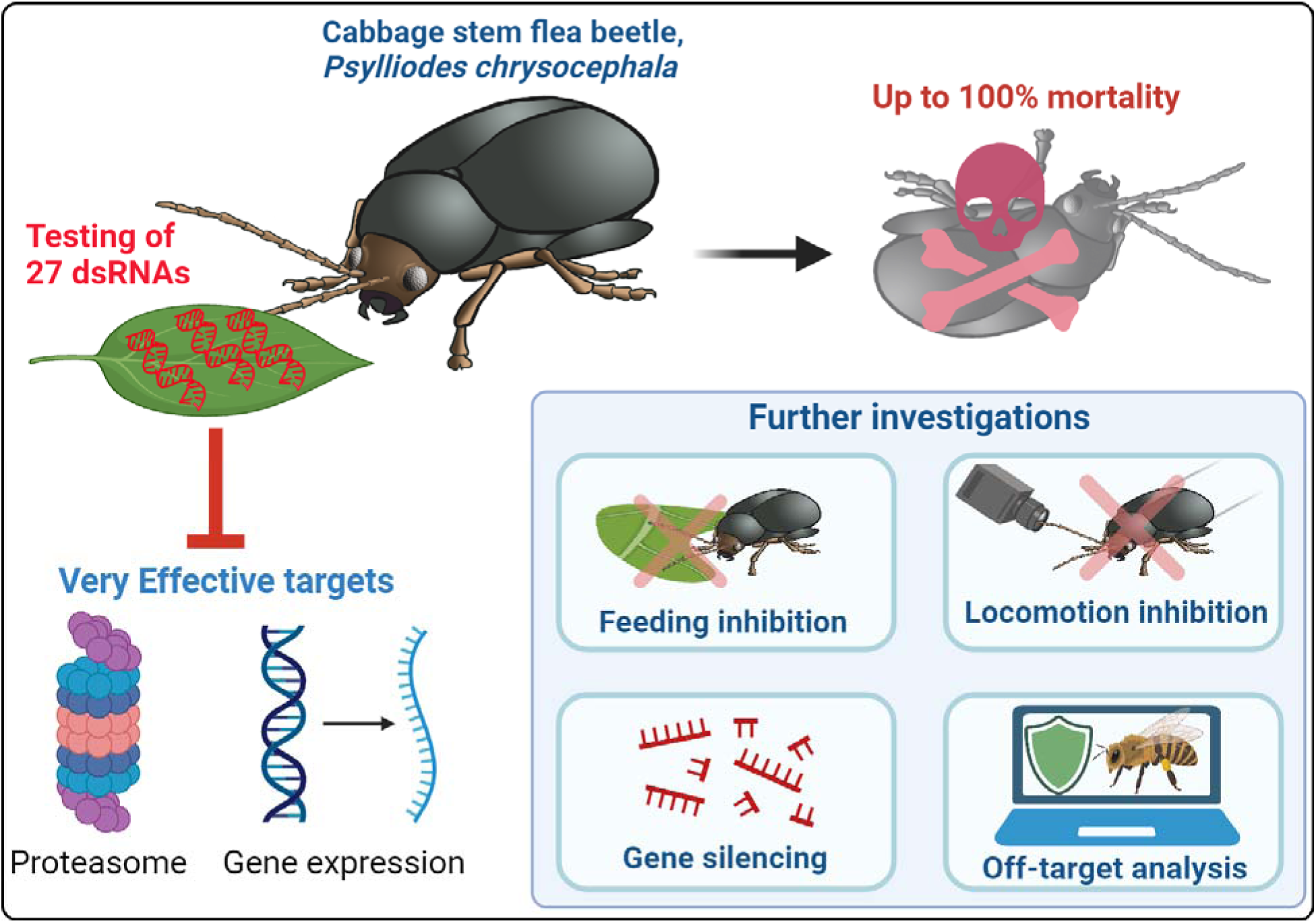

- An oral RNAi screen consisting of 27 dsRNAs identified effective target genes for RNAi-based pest control in the cabbage stem flea beetle
- The most effective target genes were involved in proteasomal pathways or gene expression, and their sequences were subjected to *in silico* off-target analysis
- Knock-down of some effective target genes also inhibited feeding or locomotion

## References

Bachman, P.M., Bolognesi, R., Moar, W.J., Mueller, G.M., Paradise, M.S., Ramaseshadri, P., Tan, J., Uffman, J.P., Warren, J., Wiggins, B.E. & Levine, S.L. (2013) Characterization of the spectrum of insecticidal activity of a double-stranded RNA with targeted activity against Western Corn Rootworm (*Diabrotica virgifera virgifera* LeConte). Transgenic Research, 22(6), 1207–1222. Available online: 10.1007/s11248-013-9716-5.

Baum, J.A., Bogaert, T., Clinton, W., Heck, G.R., Feldmann, P., Ilagan, O., Johnson, S., Plaetinck, G., Munyikwa, T., Pleau, M., Vaughn, T. & Roberts, J. (2007) Control of coleopteran insect pests through RNA interference. Nature Biotechnology, 25(11), 1322–1326. Available online: 10.1038/nbt1359.

Bolognesi, R., Ramaseshadri, P., Anderson, J., Bachman, P., Clinton, W., Flannagan, R., Ilagan, O., Lawrence, C., Levine, S., Moar, W., Mueller, G., Tan, J., Uffman, J., Wiggins, E., Heck, G. & Segers, G. (2012) Characterizing the Mechanism of Action of Double-Stranded RNA Activity against Western Corn Rootworm (*Diabrotica virgifera virgifera* LeConte). PLOS ONE, 7(10), e47534. Available online: 10.1371/journal.pone.0047534.

Buer, B., Dönitz, J., Milner, M., Mehlhorn, S., Hinners, C., Siemanowski-Hrach, J., Ulrich, J.K., Großmann, D., Cedden, D., Nauen, R., Geibel, S. & Bucher, G. (2024) Superior target genes and pathways for RNAi mediated pest control revealed by genome wide analysis in the red flour beetle *Tribolium castaneum*. bioRxiv, 2024.01.24.577003. Available online: 10.1101/2024.01.24.577003.

Cedden, D., Güney, G., Scholten, S. & Rostás, M. (2024) Lethal and sublethal effects of orally delivered double-stranded RNA on the cabbage stem flea beetle, *Psylliodes chrysocephala*. Pest Management Science, 80(5), 2282–2293. Available online: 10.1002/ps.7494.

Cheatle Jarvela, A.M., Trelstad, C.S. & Pick, L. (2020) Regulatory gene function handoff allows essential gene loss in mosquitoes. Communications Biology, 3(1), 1–10. Available online: 10.1038/s42003-020-01203-w.

Chen, J., Peng, Y., Zhang, H., Wang, K., Tang, Y., Gao, J., Zhao, C., Zhu, G., Palli, S.R. & Han, Z. (2021) Transcript level is a key factor affecting RNAi efficiency. Pesticide Biochemistry and Physiology, 176, 104872. Available online: 10.1016/j.pestbp.2021.104872.

Deutsch, C.A., Tewksbury, J.J., Tigchelaar, M., Battisti, D.S., Merrill, S.C., Huey, R.B. & Naylor, R.L. (2018) Increase in crop losses to insect pests in a warming climate. Science, 361(6405), 916–919. Available online: 10.1126/science.aat3466.

van Dijk, M., Morley, T., Rau, M.L. & Saghai, Y. (2021) A meta-analysis of projected global food demand and population at risk of hunger for the period 2010–2050. Nature Food, 2(7), 494–501. Available online: 10.1038/s43016-021-00322-9.

Emms, D.M. & Kelly, S. (2019) OrthoFinder: phylogenetic orthology inference for comparative genomics. Genome Biology, 20(1), 238. Available online: 10.1186/s13059-019-1832-y.

Feyereisen, R. (1995) Molecular biology of insecticide resistance. Toxicology Letters, 82–83, 83–90. Available online: 10.1016/0378-4274(95)03470-6.

Güney, G., Cedden, D., Körnig J., Ulber, B., Beran F., Scholten S., & Rostás M. (2024) Physiological and Transcriptional Changes Associated with Obligate Aestivation in the Cabbage Stem Flea Beetle (*Psylliodes chrysocephala*). bioRxiv, April 11, 2024. 10.1101/2024.04.08.588545.

Gu, D., Andreev, K. & Dupre, M.E. (2021) Major Trends in Population Growth Around the World. China CDC Weekly, 3(28), 604–613. Available online: 10.46234/ccdcw2021.160.

Højland, D.H., Nauen, R., Foster, S.P., Williamson, M.S. & Kristensen, M. (2015) Incidence, Spread and Mechanisms of Pyrethroid Resistance in European Populations of the Cabbage Stem Flea Beetle, *Psylliodes chrysocephala* L. (Coleoptera: Chrysomelidae). PLOS ONE, 10(12), e0146045. Available online: 10.1371/journal.pone.0146045.

Kim, K., Lee, Y.S., Harris, D., Nakahara, K. & Carthew, R.W. (2006) The RNAi pathway initiated by Dicer-2 in *Drosophila*. Cold Spring Harbor Symposia on Quantitative Biology, 71, 39–44. Available online: 10.1101/sqb.2006.71.008.

Knorr, E., Billion, A., Fishilevich, E., Tenbusch, L., Frey, M.L.F., Rangasamy, M., Gandra, P., Arora, K., Lo, W., Geng, C., Vilcinskas, A. & Narva, K.E. (2021) Knockdown of Genes Involved in Transcription and Splicing Reveals Novel RNAi Targets for Pest Control. Frontiers in Agronomy, 3. Available online: https://www.frontiersin.org/articles/10.3389/fagro.2021.715823 [Accessed 17/02/2024].

Knorr, E., Fishilevich, E., Tenbusch, L., Frey, M.L.F., Rangasamy, M., Billion, A., Worden, S.E., Gandra, P., Arora, K., Lo, W., Schulenberg, G., Valverde-Garcia, P., Vilcinskas, A. & Narva, K.E. (2018) Gene silencing in *Tribolium castaneum* as a tool for the targeted identification of candidate RNAi targets in crop pests. Scientific Reports, 8(1), 2061. Available online: 10.1038/s41598-018-20416-y.

Langmead, B., Trapnell, C., Pop, M. & Salzberg, S.L. (2009) Ultrafast and memory-efficient alignment of short DNA sequences to the human genome. Genome Biology, 10(3), R25. Available online: 10.1186/gb-2009-10-3-r25.

Li, Z., Costamagna, A.C., Beran, F. & You, M. (2024) Biology, Ecology, and Management of Flea Beetles in Brassica Crops. Annual Review of Entomology, 69(1), 199–217. Available online: 10.1146/annurev-ento-033023-015753.

Liu, S., Jaouannet, M., Dempsey, D.A., Imani, J., Coustau, C. & Kogel, K.-H. (2020) RNA-based technologies for insect control in plant production. Biotechnology Advances, 39, 107463. Available online: 10.1016/j.biotechadv.2019.107463.

Liu, X., Jiang, F., Kalidas, S., Smith, D. & Liu, Q. (2006) Dicer-2 and R2D2 coordinately bind siRNA to promote assembly of the siRISC complexes. RNA, 12(8), 1514–1520. Available online: 10.1261/rna.101606.

Manni, M., Berkeley, M.R., Seppey, M., Simão, F.A. & Zdobnov, E.M. (2021) BUSCO Update: Novel and Streamlined Workflows along with Broader and Deeper Phylogenetic Coverage for Scoring of Eukaryotic, Prokaryotic, and Viral Genomes. Molecular Biology and Evolution, 38(10), 4647–4654. Available online: 10.1093/molbev/msab199.

Mao, Y.-B., Cai, W.-J., Wang, J.-W., Hong, G.-J., Tao, X.-Y., Wang, L.-J., Huang, Y.-P. & Chen, X.-Y. (2007) Silencing a cotton bollworm P450 monooxygenase gene by plant-mediated RNAi impairs larval tolerance of gossypol. Nature Biotechnology, 25(11), 1307–1313. Available online: 10.1038/nbt1352.

Marshall, R.S. & Vierstra, R.D. (2019) Dynamic Regulation of the 26S Proteasome: From Synthesis to Degradation. Frontiers in Molecular Biosciences, 6. Available online: https://www.frontiersin.org/articles/10.3389/fmolb.2019.00040 [Accessed 17/02/2024].

Martinez, J., Patkaniowska, A., Urlaub, H., Lührmann, R. & Tuschl, T. (2002) Single-Stranded Antisense siRNAs Guide Target RNA Cleavage in RNAi. Cell, 110(5), 563–574. Available online: 10.1016/S0092-8674(02)00908-X.

Mehlhorn, S., Hunnekuhl, V.S., Geibel, S., Nauen, R. & Bucher, G. (2021) Establishing RNAi for basic research and pest control and identification of the most efficient target genes for pest control: a brief guide. Frontiers in Zoology, 18(1), 60. Available online: 10.1186/s12983-021-00444-7.

Mehlhorn, S., Ulrich, J., Baden, C.U., Buer, B., Maiwald, F., Lueke, B., Geibel, S., Bucher, G. & Nauen, R. (2021) The mustard leaf beetle, *Phaedon cochleariae*, as a screening model for exogenous RNAi-based control of coleopteran pests. Pesticide Biochemistry and Physiology, 176, 104870. Available online: 10.1016/j.pestbp.2021.104870.

Naganuma, M., Tadakuma, H. & Tomari, Y. (2021) Single-molecule analysis of processive double-stranded RNA cleavage by Drosophila Dicer-2. Nat. Commun., 12. Available online: 10.1038/s41467-021-24555-1.

Oosthoek, S. (2013) Pesticides spark broad biodiversity loss. Nature. Available online: 10.1038/nature.2013.13214.

Ortega-Ramos, P.A., Coston, D.J., Seimandi-Corda, G., Mauchline, A.L. & Cook, S.M. (2022) Integrated pest management strategies for cabbage stem flea beetle (*Psylliodes chrysocephala*) in oilseed rape. GCB Bioenergy, 14(3), 267–286. Available online: 10.1111/gcbb.12918.

Pack, C.-G., Yukii, H., Toh-e, A., Kudo, T., Tsuchiya, H., Kaiho, A., Sakata, E., Murata, S., Yokosawa, H., Sako, Y., Baumeister, W., Tanaka, K. & Saeki, Y. (2014) Quantitative live-cell imaging reveals spatio-temporal dynamics and cytoplasmic assembly of the 26S proteasome. Nature Communications, 5(1), 3396. Available online: 10.1038/ncomms4396.

Parker, K.M., Barragán Borrero, V., van Leeuwen, D.M., Lever, M.A., Mateescu, B. & Sander, M. (2019) Environmental Fate of RNA Interference Pesticides: Adsorption and Degradation of Double-Stranded RNA Molecules in Agricultural Soils. Environmental Science & Technology, 53(6), 3027–3036. Available online: 10.1021/acs.est.8b05576.

Preall, J.B. & Sontheimer, E.J. (2005) RNAi: RISC Gets Loaded. Cell, 123(4), 543–545. Available online: 10.1016/j.cell.2005.11.006.

Reinders, J.D., Moar, W.J., Head, G.P., Hassan, S. & Meinke, L.J. (2023) Effects of SmartStax® and SmartStax® PRO maize on western corn rootworm (*Diabrotica virgifera virgifera* LeConte) larval feeding injury and adult life history parameters. PLOS ONE, 18(7), e0288372. Available online: 10.1371/journal.pone.0288372.

Rodrigues, T.B., Mishra, S.K., Sridharan, K., Barnes, E.R., Alyokhin, A., Tuttle, R., Kokulapalan, W., Garby, D., Skizim, N.J., Tang, Y., Manley, B., Aulisa, L., Flannagan, R.D., Cobb, C. & Narva, K.E. (2021) First Sprayable Double-Stranded RNA-Based Biopesticide Product Targets Proteasome Subunit Beta Type-5 in Colorado Potato Beetle (*Leptinotarsa decemlineata*). Frontiers in Plant Science, 12. Available online: https://www.frontiersin.org/article/10.3389/fpls.2021.728652.

Sivčev, L., Graora, D., Sivčev, I., Tomić, V. & Dudić, B. (2016) Phenology of cabbage stem flea beetle (*Psylliodes chrysocephala* L.) in oilseed rape. Pesticidi i Fitomedicina, 31(3–4), 139–144.

Sparks, T.C., Crossthwaite, A.J., Nauen, R., Banba, S., Cordova, D., Earley, F., Ebbinghaus-Kintscher, U., Fujioka, S., Hirao, A., Karmon, D., Kennedy, R., Nakao, T., Popham, H.J.R., Salgado, V., Watson, G.B., Wedel, B.J. & Wessels, F.J. (2020) Insecticides, biologics and nematicides: Updates to IRAC’s mode of action classification - a tool for resistance management. Pesticide Biochemistry and Physiology, 167, 104587. Available online: 10.1016/j.pestbp.2020.104587.

Sparks, T.C. & Nauen, R. (2015) IRAC: Mode of action classification and insecticide resistance management. Pesticide Biochemistry and Physiology, 121, 122–128. Available online: 10.1016/j.pestbp.2014.11.014.

Tanaka, K. & Ichihara, A. (1989) Half-life of proteasomes (multiprotease complexes) in rat liver. Biochemical and Biophysical Research Communications, 159(3), 1309–1315. Available online: 10.1016/0006-291X(89)92253-5.

Ulrich, J., Dao, V.A., Majumdar, U., Schmitt-Engel, C., Schwirz, J., Schultheis, D., Ströhlein, N., Troelenberg, N., Grossmann, D., Richter, T., Dönitz, J., Gerischer, L., Leboulle, G., Vilcinskas, A., Stanke, M. & Bucher, G. (2015) Large scale RNAi screen in *Tribolium* reveals novel target genes for pest control and the proteasome as prime target. BMC Genomics, 16(1), 674. Available online: 10.1186/s12864-015-1880-y.

Willis, C.E., Foster, S.P., Zimmer, C.T., Elias, J., Chang, X., Field, L.M., Williamson, M.S. & Davies, T.G.E. (2020) Investigating the status of pyrethroid resistance in UK populations of the cabbage stem flea beetle (*Psylliodes chrysocephala*). Crop Protection, 138, 105316. Available online: 10.1016/j.cropro.2020.105316.

Willow, J. & Veromann, E. (2021) Highly Variable Dietary RNAi Sensitivity Among Coleoptera. Frontiers in Plant Science, 12. Available online: https://www.frontiersin.org/article/10.3389/fpls.2021.790816.

Zeisel, A., Köstler, W.J., Molotski, N., Tsai, J.M., Krauthgamer, R., Jacob-Hirsch, J., Rechavi, G., Soen, Y., Jung, S., Yarden, Y. & Domany, E. (2011) Coupled pre-mRNA and mRNA dynamics unveil operational strategies underlying transcriptional responses to stimuli. Molecular Systems Biology, 7, 529. Available online: 10.1038/msb.2011.62.

Zhang, J., Khan, S.A., Hasse, C., Ruf, S., Heckel, D.G. & Bock, R. (2015) Full crop protection from an insect pest by expression of long double-stranded RNAs in plastids. Science, 347(6225), 991–994. Available online: 10.1126/science.1261680.

