## Supporting information for "Effective target genes for RNA interference-based management of the cabbage stem flea beetle"

Supporting Table 1. List of 14 effective lethal dsRNAs at 500 ng, annotations of the respective targeted genes, NCBI accessions (TSA: GKIH01000001 : GKIH01098921), and the log-rank survival analysis results

| dsRNA | Annotation^1^ | ID / Accession | % Mortality in 14 d | Median survival (d) | Hazard ratio^2^ | *P*^2^ | Gene list in Buer et al., (2024)^3^ |
| --- | --- | --- | --- | --- | --- | --- | --- |
| dsRpt4 | 26S proteasome regulatory subunit 10B | TRINITY_DN5454_c0_g1  GKIH01032540.1 | 100 | 8 | 19.8 | <.001 | Cluster 1  1 transfer |
| dsProsβ6 | Proteasome subunit beta type-1 | TRINITY_DN20894_c0_g1  GKIH01084533.1 | 100 | 8 | 19.4 | <.001 | Cluster 1  1 transfer |
| dsRpt1 | 26S proteasome regulatory subunit 7 | TRINITY_DN15123_c0_g1  GKIH01096120.1 | 100 | 9 | 19.1 | <.001 | Cluster 1  1 transfer |
| dsSap30 | SAP30-binding | TRINITY_DN1168_c0_g1  GKIH01077053.1 | 100 | 8 | 24.3 | <.001 | Cluster 1  no transfer |
| dsRpt3 | 26S proteasome regulatory subunit 6B | TRINITY_DN5454_c0_g3  GKIH01032542.1 | 89.5 | 9 | 18.2 | <.001 | Cluster 1  1 transfer |
| dsPAbp | Polyadenylate-binding 1 Short=PABP-1 Short=Poly(A)-binding 1 | TRINITY_DN46_c1_g1  GKIH01077302.1 | 85 | 8 | 17.9 | <.001 | Cluster 1  2 transfers |
| dsRpn7 | 26S proteasome non-ATPase regulatory subunit 6 | TRINITY_DN85218_c0_g1  GKIH01075228.1 | 85 | 8 | 18.4 | <.001 | Cluster 1  2 transfers |
| dsβCOP | Coatomer subunit beta | TRINITY_DN6946_c0_g1  GKIH01041308.1 | 82.8 | 8 | 19.1 | <.001 | Cluster 1  1 transfer |
| dsProsβ7 | Proteasome subunit beta type-4 | TRINITY_DN6590_c0_g1  GKIH01027377.1 | 81.25 | 8 | 16.67 | <.001 | Cluster 1  2 transfers |
| dsSf3b1 | Splicing factor 3B subunit 1 | TRINITY_DN2158_c0_g2  GKIH01026338.1 | 73.2 | 10 | 13.5 | <.001 | Cluster 1  1 transfer |
| dsRpL11 | 60S ribosomal L11 | TRINITY_DN93688_c0_g1  GKIH01029471.1 | 70 | 13 | 14.9 | <.001 | Cluster 1  no transfer |
| dsChc | Clathrin heavy chain | TRINITY_DN679_c0_g1  GKIH01054170.1 | 62.5 | 11 | 12.3 | <.001 | Cluster 1  1 transfer |
| dsRpt2 | 26S proteasome regulatory subunit 4 Short=P26s4 | TRINITY_DN4712_c0_g1  GKIH01096759.1 | 60 | 11.5 | 11.3 | <.001 | Cluster 1  1 transfer |
| dsRpL1Ab | 60S ribosomal L10a | TRINITY_DN1232_c0_g1  GKIH01013468.1 | 50 | 14 | 9.59 | <.001 | Cluster 1  no transfers |

^1^ The annotations of the CSFB orthologs were taken from *Tribolium castaneum* annotations in Buer et al., (2024)

^2^ The hazard ratios and P values were obtained by comparing the survival curves of the lethal dsRNA treatment with that of dsmGFP control via log-rank test. All P values were significant after Bonferroni correction for the 13 pairwise tests.

^3^ Buer et al., (2024) presented Clusters 1-5 based on dose-response of effective target genes in *T. castaneum*. A subset of these were successfully transferred to only *Phaedon cochleariae* (1 transfer) or both *P. cochleariae* and *Leptinotaras decemlineata* (2 transfers)

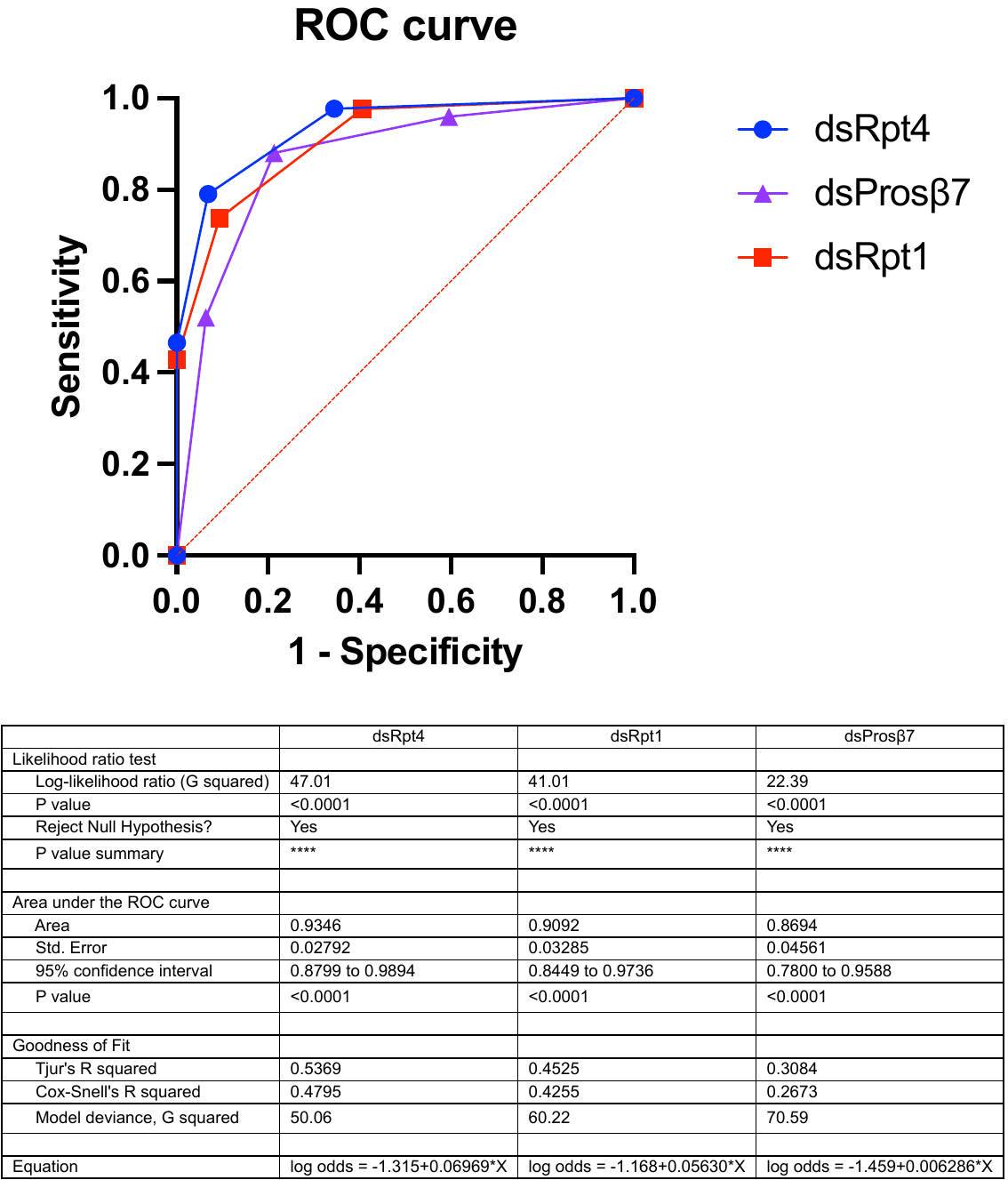

Supporting Figure 1. The ROC (receiver operating characteristic) curves of the logistic regression analysis to investigate the dose responses to dsRpt4, dsProsβ7, and dsRpt1. The table gives statistics regarding the performance of the models (calculated using GraphPad v10.2).

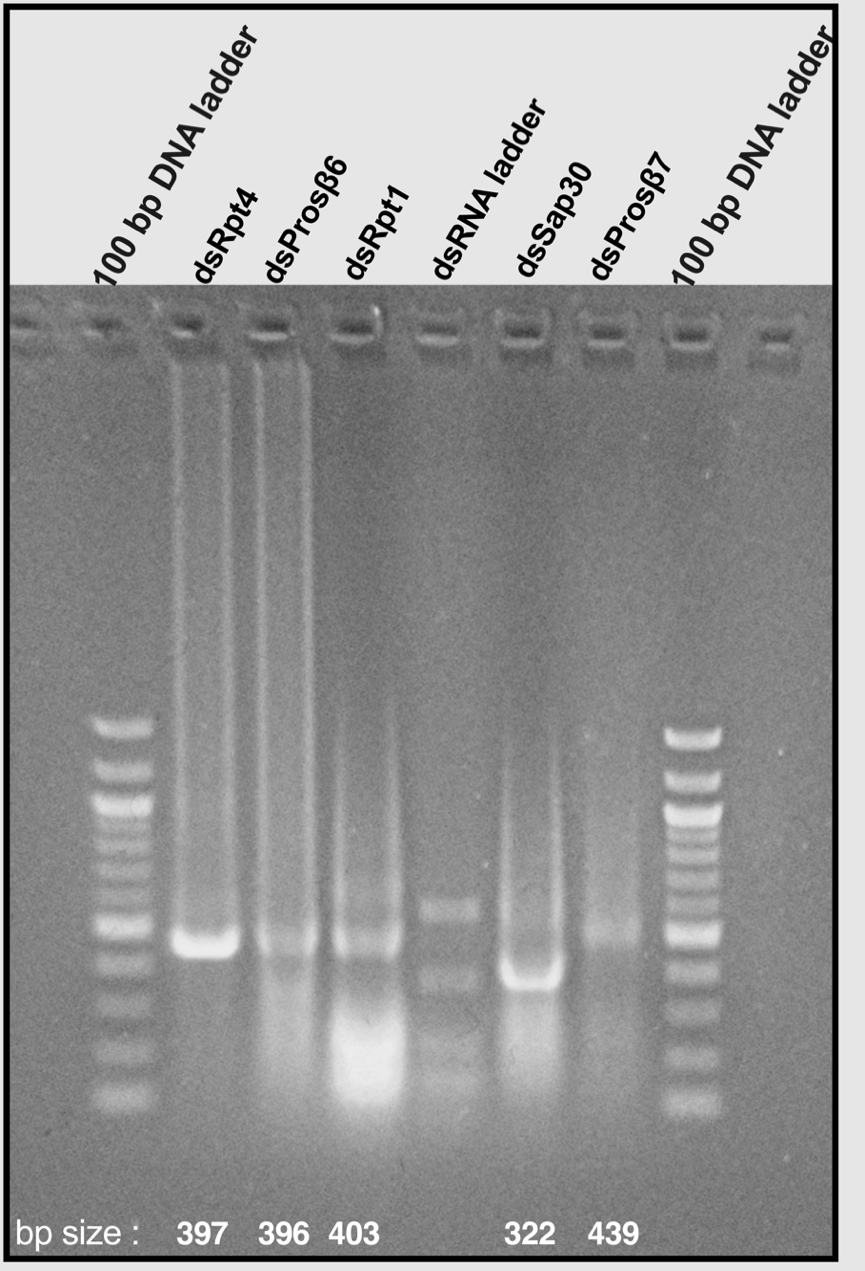

Supporting Figure 2. Example of visualization of double-stranded RNAs on 1.5% agarose gel. Two µL of dsRNA Ladder (NEB Catalog # N0363S) was run in parallel as a reference. The sizes of the dsRNA ladders are 500, 300, 150, 80, 30, and 21 base pairs. The dsRNAs were loaded at 2.5 µg per column. The expected base pair sizes of the dsRNAs based on the length of the amplicons targeted by the PCR primers are given below the bands. A contrast adjustment was applied uniformly.

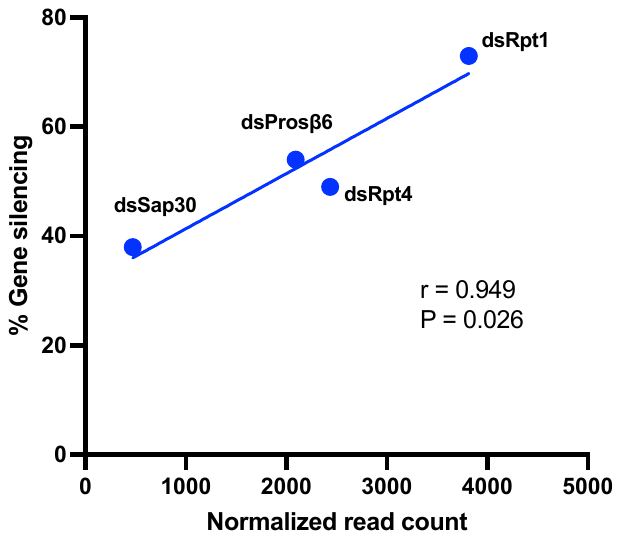

Supporting Figure 3. Pearson correlation analysis between the achieved percent gene silencing through the oral delivery of 500 ng dsRNA and the basal expression (median transcripts per million data from RNA-seq on females with n = 7) of the respective gene in adult cabbage stem flea beetle.
